# Adenoviral-mediated gene transfer into the scala media of the mouse cochlea in vivo

**DOI:** 10.64898/2026.04.25.720849

**Authors:** Fukuichiro Iguchi, Debbie Bratt, Ming Xiao, Amy D. Erdman, Amanda Sekijima, Clifford R. Hume

## Abstract

Gene therapy may provide a way to restore inner ear function to deaf and dizzy patients. The mouse is a crucial model system for functional genomics because of the numerous genetic models for hearing loss and inner ear dysfunction. Using an advance generation, E1-/E3-/E2b-(preterminal protein-/polymerase-) Type 5 Adenovirus, we investigated several routes of virus microinjection to determine which were most reproducible in targeting the endolymphatic fluid compartment of the cochlea. We found that when adenovirus is injected via the round window, transduced cells are found only adjacent to the scala tympani and not in the organ of Corti, suggesting that Adenovirus is unable to penetrate the basilar membrane or bony wall of the modiolus. Delivery to the cochlea via the semicircular canals is also inefficient. In contrast, our new method, via a stylomastoid foramen cochleostomy, increases the likelihood of adenovirus gene transfer to the scala media including cells in the organ of Corti and stria vascularis while preserving some hearing. The ability to target delivery of virus and other therapeutic reagents to specific inner ear fluid compartments will facilitate in vivo testing of candidate molecules implicated in multiple aspects of inner ear physiology and regeneration.

## 1. Introduction

Gene therapy may provide a way to restore function to patients with hearing and balance disorders. Experimental gene delivery to the inner ear in mammals was first reported in 1996 (Raphael et al., 1996). Since then there has been dramatic progress in the understanding of the genetics of hearing loss, identification of candidate therapeutic genes and techniques for delivery to the inner ear. More than 100 loci have been mapped that can lead to non-syndromic hearing loss (Van Camp and Smith; Friedman and Griffith, 2003). Many of the gene products altered or deficient in these individuals have also been identified. In addition to the discovery of mutations responsible for hearing loss, many of the critical molecules that regulate inner ear embryonic development have been identified (Barald and Kelley, 2004).

Many of the inner ear gene therapy studies that have shown promise were conducted in guinea pig (Luebke et al., 2001; Luebke et al., 2001; Kawamoto et al., 2003; Nakaizumi et al., 2004; Izumikawa et al., 2005; Luebke et al., 2009; Shibata et al., 2010; Wise et al., 2010). This species has the advantage of a large inner ear allowing direct microinjection into the scalae and targeting of specific cell types, but lacks genetic models of hearing loss. Because of the small size of the cochlea, reproducible, atraumatic delivery of gene therapy vectors to the mouse inner ear has been challenging (Jero et al., 2001; Liu et al., 2005; Liu et al., 2007; Staecker et al., 2007; Wenzel et al., 2007; Baker et al., 2009). On the other hand, mice have the huge advantage of numerous genetic models for hearing loss and inner ear dysfunction, more rapid breeding and less husbandry expense. Recently, a number of deafening protocols have also been developed to study acquired hearing loss in mice (Kujawa and Liberman, 2006; Oesterle et al., 2008; Kujawa and Liberman, 2009; Oesterle and Campbell, 2009). These similarities in structure, physiology and genetics make the mouse a crucial model system for functional genomics of the auditory system (Brown et al., 2008).

We investigated whether alternate routes of virus microinjection into the mouse inner ear were more reproducible in targeting the endolymphatic fluid compartment, the scala media, because it contains the major sensory and functional components of the inner ear. Based on published studies, the most promising surgical approaches to deliver chemicals, viruses or cells atraumatically to the inner ear and preserve hearing are via a semicircular canalostomy, the ampulla, basal turn cochleostomy and round window membrane (Lalwani et al., 1996; Raphael et al., 1996; Praetorius et al., 2003; Iguchi et al., 2004; Chen et al., 2006; Wenzel et al., 2007). We describe a method to deliver adenoviral vectors to the scala media of the mouse cochlea with some preservation of hearing (loss of ∼ 35 dB). With this technique, the primary sites of transduction are supporting cells in the organ of Corti, Reissner’s membrane and the stria vascularis. We believe that this method will be valuable for in vivo screening of candidate genes implicated in inner ear function and regeneration in mice.

## 2. Materials and Methods

### 2.1. Virus Preparation

#### Adenovirus Production

Initial studies using early generation E1/E3 deleted adenovirus showed evidence of damage to hair cell stereocilia in vitro and loss of hearing (Holt et al., 1999; Luebke et al., 2001; Luebke et al., 2001). While some of this damage may be due to surgical trauma, more extensively deleted adenoviral vectors lacking either the E4 region or viral polymerase, or Adeno-associated virus appear to be less toxic to the inner ear (Holt et al., 1999; Luebke et al., 2001; Kawamoto et al., 2003; Corey et al., 2004; Liu et al., 2005; Wenzel et al., 2007). This suggests that residual adenoviral gene expression may be directly or indirectly (through an immune response) responsible for subsequent toxicity or hearing loss. To minimize the potential effects of toxicity related to adenoviral gene expression, we used a later generation Type 5 Adenovirus deleted for polymerase (E1-/E3-/pol-), or preterminal protein and polymerase (E1-/E3-/TP-/pol-) that has been shown to cause minimal direct toxicity to the inner ear (Amalfitano and Chamberlain, 1997; Holt et al., 1999; Luebke et al., 2001; Luebke et al., 2001; Holt, 2002). To identify targeted cells, we used a Green Fluorescent Protein (hrGFP or EGFP) reporter gene driven by the human cytomegalovirus (CMV) promoter. Replication incompetent adenovirus was produced using the AdEasy system with the more extensively deleted AdEasy backbone vector, pAdE-(delta)TP(delta)pol or pAdE-(delta)pol and pShuttle-CMV-IRES-hrGFP2 or pShuttle CMV-EGFP (Stratagene) (Amalfitano and Chamberlain, 1997; Wise et al., 2010). Appropriately recombined viral DNA’s were transfected into C7-HEK 293 cells that stably expresses Adenovirus polymerase and pre-terminal protein in addition to E1 functions. Crude virus (AdE-CMV-GFP, AdE-CMV-hrGFP) was amplified through several rounds to produce a high titer and purified using commercially available ion exchange chromatography membranes (Vivascience, Sartorius). The purified virus was titered by spectrophotometry (>1 × 10^12^ gc/ml by OD_260_) and on cultured cell lines (HT-1080) to verify GFP expression. Titers on HT 1080 cells ranged from 20-50% of OD_260_. All viral stocks were verified to be free of replication competent Adenovirus using a semi-quantitative PCR assay for E1a and E1b (Suzuki et al., 2004). Virus stocks were aliquoted and stored at -80°C before use.

Immediately before a series of injections, a small quantity of the virus stock was diluted (10:4:6) with 5x concentrated artificial endolymph (1x stock= 140 mM KCl, 25mM KHCO_3_) (Salt and DeMott, 1997) and blue food dye to allow real-time monitoring of injections (FD&C Blue 1, 0.1% propylparaben, propylene glycol, McCormick/Shilling)(Peng et al., 2009). The University of Washington Environmental Health and Safety Institutional Biosafety Committee approved all procedures.

### 2.2. Surgical Procedure

Adult Swiss Webster mice weighing between 20 and 30 g were used for the study. The University of Washington Institutional Animal Care and Use Committee approved all procedures.

For virus injections, the mice were anesthetized using ketamine hydrochloride (130 mg/kg, i.p.) and xylazine hydrochloride (8.8 mg/kg, i.p.). All procedures were performed on the right ear. A postauricular incision was made and the tympanic bulla and the facial nerve were exposed. The facial nerve was followed upward in order to identify the stylomastoid foramen. For all injections, the presence of a blue dye in the fluid allowed direct monitoring of both the progress of delivery to the labyrinth and leakage from the injection site. In each paradigm, animals with gross leakage of blue dye after injection were sacrificed and excluded from further analysis. In some cases, Skin Shield Liquid Bandage (Del Pharmaceuticals, NY, USA) was used to minimize leakage around the injection needle (A. N. Salt, ARO 2009).

#### Posterior semicircular canal (PSC) injection

We followed the same technique that was used to deliver neomycin to the inner ear in a previous report (Nakagawa et al., 2003; Iguchi et al., 2004). The lateral semicircular canal is located dorsal to the stylomastoid foramen. The PSC is identified at the posterior edge of the temporal bone. A small hole (about 180 μm in diameter) was made in each of the semicircular canals using a 26-gauge needle. A blunt fused silica needle was inserted into the PSC. Three microliters of virus suspension (approximately 10^9^ viral particles by OD) or virus storage buffer was slowly injected by hand over three minutes. The holes in the semicircular canals were plugged with the connective tissue. Total operating time was approximately 10 minutes.

#### Round window (RW) injection

The upper posterior corner of the tympanic bulla was perforated to visualize the round window. After drying the round window membrane, Skin Shield Liquid Bandage (Del Pharmaceuticals, NY, USA) was applied to the round window membrane in several layers, allowing it to dry between each layer. A glass micropipette with an outer tip diameter of approximately 10 μm was loaded with virus suspension or virus storage buffer and the tip slowly advanced through the round window membrane into the scala tympani using a micromanipulator (Narishige, Tokyo, Japan). Injections of 0.2 μl were performed using a computer controlled micro pump (Ultra Micro Pump III, World Precision Instruments, FL) at a rate of 0.1 μl/min. After the injection of fluid, the hole was sealed with additional layers of liquid bandage to seal the hole. Total operating time was approximately 20 minutes.

#### Stylomastoid foramen (SF) injection

This approach is shown in Figure 1. For illustrative purposes, the bulla and external auditory canal were widely opened to demonstrate the relevant anatomy. A more limited exposure was utilized in experimental animals. The bone in the bottom of the stylomastoid foramen niche was carefully picked away using a 28G needle until the lateral wall of the cochlear duct was revealed. The diameter of the hole was about 100 μm. After verifying that the cochlear duct was intact and that there was no endolymph leakage, liquid bandage was applied to the hole in several thin layers as above. A pulled glass micropipette loaded with virus suspension or virus storage buffer was slowly advanced through the stria vascularis into the scala media using the same equipment as the RW injection. The 0.2 μl of virus suspension, or loading buffer was injected at the same rate as above. After the injection, the hole was sealed with several additional layers of liquid bandage. Total operating time was approximately 30 minutes.

**Figure 1.**
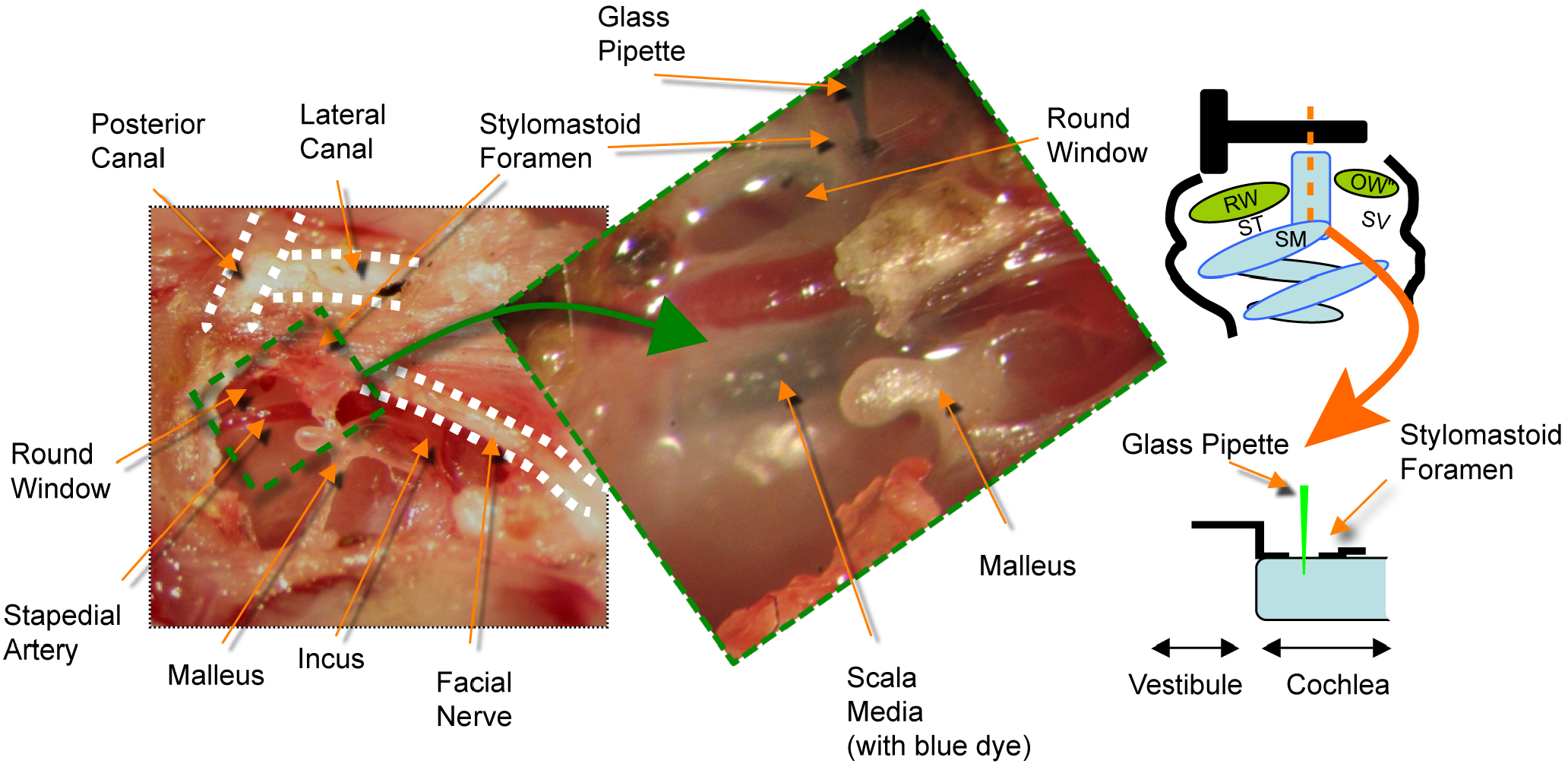
Surgical anatomy of the stylomastoid foramen (SF) technique. The orange dashed line in the schematic denotes the trajectory of the injection needle to the endolymphatic fluid compartment (blue) and the scala media. For illustration purposes, the bulla and external auditory canal are widely opened to show the relevant anatomy. RW= round window, OW= oval window, ST= scala tympani, SV= scala vestibuli, SM= scala media.

### 2.3. Auditory Function Measurements

To assess the impact of the injection procedure on the physiological status of experimental ears, we measured auditory brainstem response (ABR) thresholds in the right ear. ABR measurements at 4, 8, 16 and 32 kHz pure tones were obtained prior to surgery and seven days after surgery in each group.

Mice were anesthetized by intraperitoneal administration of xylazine (8.8 mg/kg; Bayer, Shawnee Mission, KS) and ketamine-HCl (130 mg/kg; Parke Davis, Fort Dodge, IA) and placed in a sound-attenuating chamber (IAC, New York, NY, USA). To ensure consistent results, the body temperature of each animal was maintained at 36–38 °C, using an isotherm heating pad.

Sound stimuli were generated digitally, attenuated using programmable attenuators, and delivered via electrostatic speakers (RP-2, PA-5, ED-1, and EC-1; Tucker-Davis Technologies, Alachua, FL, USA) connected to the ear by a plastic tube inserted into the external canal. The stimuli were 4 ms at a rate of 20/s (1 ms rise/fall, 2 ms plateau, 50-ms intervals). Prior to the ABR recording, the speakers were calibrated across the frequency range using a probe tube microphone (Type 4133; Bruel & Kjaer, Denmark).

ABR recordings were collected from subdermal electrodes placed at the vertex (positive), ipsilateral mastoid (negative), and thigh (ground). The signals were amplified 10,000 times using a biological amplifier (HS4/DB4; Tucker-Davis Technologies) and filtered via a two-step process. Initial amplification of 1000 times was made using a preamplifier with a 10-to 10,000-Hz filter (DAM-6A Differential Pre-amp, World Precision Instruments, Sarasota, FL, USA) and a secondary amplification was made using an amplification of 100 times and a 0.3-to 3-kHz filter by the TDT System II spike conditioner.

500 trials were averaged at each frequency with tone bursts in systematic 5-dB steps. The threshold was defined as the lowest level at which waves of the ABR could be clearly detected by visual inspection. If the hearing threshold was over 95 dB, then it was assigned to 100 dB.

Data were statistically analyzed using 2-way ANOVA performed with GraphPad Prism (GraphPad Software, Inc., CA, USA). As noted in the text, values of p below 0.05 were considered significant.

### 2.4. Histological Analysis

#### 2.4.1 Tissue preparation

Mice were sacrificed and the both cochleae were harvested seven days after surgery. After removal of the bulla, the stapes was lifted from the oval window and small openings were made in the apical and basilar turns. Cold 4% paraformaldehyde in 0.1M phosphate buffer, pH 7.4, was perfused slowly through the cochlea and the temporal bones fixed overnight at 4 °C. Following fixation, the temporal bones were washed three times (10 min. each) in PBS, (pH 7.4). The tissue was prepared as whole-mount preparations or cryostat sections (Hume et al., 2007; Oesterle et al., 2008). For the former, segments (half turns) of the organ of Corti were carefully dissected free from the cochlea, the stria vascularis was pulled off or trimmed down, and the tectorial membrane lifted free with fine forceps. For cryosections, the temporal bones of adult mice were decalcified in 0.4M EDTA (pH 8.0) for two days at 4 °C, rinsed in PBS, and sequentially infiltrated with sucrose (10% for 30 min., 15% for 30 min., 30% overnight, 1:1 mix 30% sucrose/O.C.T overnight) and O.C.T. compound (Tissue Tek, Sakura Finetek Inc., Torrance CA), frozen rapidly with dry ice, and cut into 10 μm sections with a cryostat. All sections were collected and were mounted onto an alternating series of 10 Platinum Superfrost (+) slides (Mercedes Medical). In contrast to some reports in guinea pig, in over 100 injected mice that we analyzed, no GFP expression was detected in the non-injected contralateral ear (Kho et al., 2000).

For relative comparisons of percentage of GFP positive cells found within the endolymphatic and perilymphatic fluid compartments of the cochlea and the vestibular region by site of injection, at least three separate slides containing serial sections (see above) from each injected ear were reviewed and tallied for the location of positive cells. Ears with no positive cells in the labyrinth (i.e., unsuccessful injections) were not considered. While this method is not strictly quantitative, it does give an estimate of “targeting” efficiency to the cochlea.

#### 2.4.2 Antibodies

The following antibodies were used to label cell populations in the inner ear: Hair cells: a mouse monoclonal anti-parvalbumin antibody (parv19, Cat. No. P3088, Sigma, St. Louis MI, 1:2000) (Sage et al., 2000), a rabbit anti-pig myosin 6 polyclonal antibody (from Dr. Tama Hasson, UC-San Diego, or Proteus Biosciences, Ramona CA, Cat. No. 25-6791, 1:1500) and a rabbit anti-human myosin 7a (from Dr. Tama Hasson or Proteus Biosciences, Cat. No. 25-6790, 1:2000) (Hasson and Mooseker, 1994; Hasson et al., 1995; Hasson et al., 1997; Hasson et al., 1997). Outer hair cells: goat anti-prestin (Santa

Cruz Biochemicals, sc22692, 1:750) (Belyantseva et al., 2000; McLean et al., 2009); Neurons: chicken anti-neurofilament (Chemicon/Millipore, AB5539, 1:5000), neurons and hair cells (rabbit anti-calretinin, Chemicon/Millipore, AB149, 1:1000). In some cases, the GFP/hrGFP signal was amplified using the appropriate antibodies (rabbit anti-hrGFP, Stratagene, 240142, 1:1000; rabbit anti-GFP, Molecular Probes/Invitrogen, A-11122, 1:500, Alexa-488 conjugated rabbit anti-GFP, Molecular Probes/Invitrogen A-21311); Supporting cells: Sox2 (rabbit anti-human Sox2, Chemicon/Millipore AB5603, 1:1000, goat anti human-Sox2 Santa Cruz Y-17/SC-17320, 1:500). The specificity of these antibodies has been verified in previous published work (Ferri et al., 2004; Kiernan et al., 2005; Hume et al., 2007; Oesterle et al., 2008)

#### 2.4.3 Immunofluorescent labeling

The tissue was permeabilized for 30 minutes with 0.1% saponin/0.1% Tween 20 in PBS. To prevent nonspecific binding of the primary antibody, tissues were then incubated for 1 hour in a blocking solution consisting of 10% normal serum/0.03% saponin/0.1% Triton X-100 in PBS. Primary antibody incubations were performed for 1 day at 4°C in PBS, 0.03% saponin, 3% serum, 2 mg/ml bovine serum albumin, and 0.1% Triton x-100 (Hume et al., 2007). Fluorescent-conjugated secondary antibodies (Alexa 488, 568, 594, 647, Invitrogen/Molecular Probes) were used at a dilution of 1:500-1:2000 in the same buffer for 2-4 hours (at room temperature) or overnight (at 4°C). Sections were washed after each antibody incubation (3 times for 10-15 min. each) in 0.1% Tween 20 in PBS. After counterstaining nuclei with bisbenzimide (Cat. No. H1398, Molecular Probes, Eugene OR, 0.5 micrograms/ml), DAPI (Cat. No. D9542, Sigma-Aldrich, 1 microgram/ml), propidium iodide (PI, Cat. No. P1304, Invitrogen Corp., 1 microgram/ml), or YO-PRO-1 (Cat. No. Y3603, Invitrogen Corp., Eugene OR, 1:200) specimens were mounted in Vectashield (Vector Laboratories), coverslipped, and examined with either epifluorescence or confocal fluorescence microscopy. At least three individual animals representative of each experimental paradigm were analyzed for each antibody.

#### 2.4.4 Microscopic imaging

Temporal bone frozen sections and whole-mount organ of Corti preparations were viewed under epifluorescence on a Zeiss Axioplan microscope, and images were captured with a Photometrics CoolSNAP hq digital camera (Image Processing Solutions, North Reading MA). Slidebook 4.0.1.40 (Intelligent Imaging Innovations, Denver CO) acquisition and processing software was used to acquire digital images and convert them to TIFF files. In some cases, whole-mount preparations and cryostat sections were viewed with an Olympus FV-1000 laser scanning confocal microscope. (405 nm blue diode, 457 nm, 488 nm and 514 nm multi-line argon, 543 nm helium neon green, 637 nm helium neon red lasers). Fluoview version 1.4a acquisition software was used. Files were imported into ImageJ 1.42a (NIH) and/or Adobe Photoshop CS version 8 (Adobe, Seattle WA) for processing and analysis.

## 3. Results

Review of the published studies on delivery of virus to the inner ear in vivo shows that targeting of gene expression is inconsistent. While some published studies using adenovirus and adeno-associated virus show widespread viral gene expression, others show more restricted distribution. Similarly, some studies *in vitro* show expression predominantly in hair cells, while others show dominant expression in supporting cells. These differences may relate to a number of variables including: the family or serotype of virus, the tissue specificity of the promoter and route of delivery. In this study we assessed the effects of site of virus delivery on gene expression and hearing. Using a high titer preparation of replication defective Type 5 Adenovirus lacking E1, E3, preterminal protein and adenoviral polymerase (AdE-CMV-GFP, see Methods), we systematically compared delivery via the Posterior Semicircular Canal (PSC), Round Window membrane (RW) and a novel cochleostomy approach to the scala media via the Stylomastoid Foramen (SF) (Figure 1).

To identify targeted cells, we used a GFP reporter gene driven by a well-characterized human cytomegalovirus (CMV) promoter that is expected to be relatively promiscuous. GFP expression detected by direct fluorescence and by indirect labeling with antibodies to GFP was comparable in all cases. Hearing was tested by ABR in mice before injection, and at the time of sacrifice one week following injection by the various routes. To control for damage caused by the injection independent from virus exposure, we also tested the hearing of animals injected with virus loading buffer. The location of GFP-expressing cells in each group of animals was assessed by review of serial sections (Figure 2).

**Figure 2.**
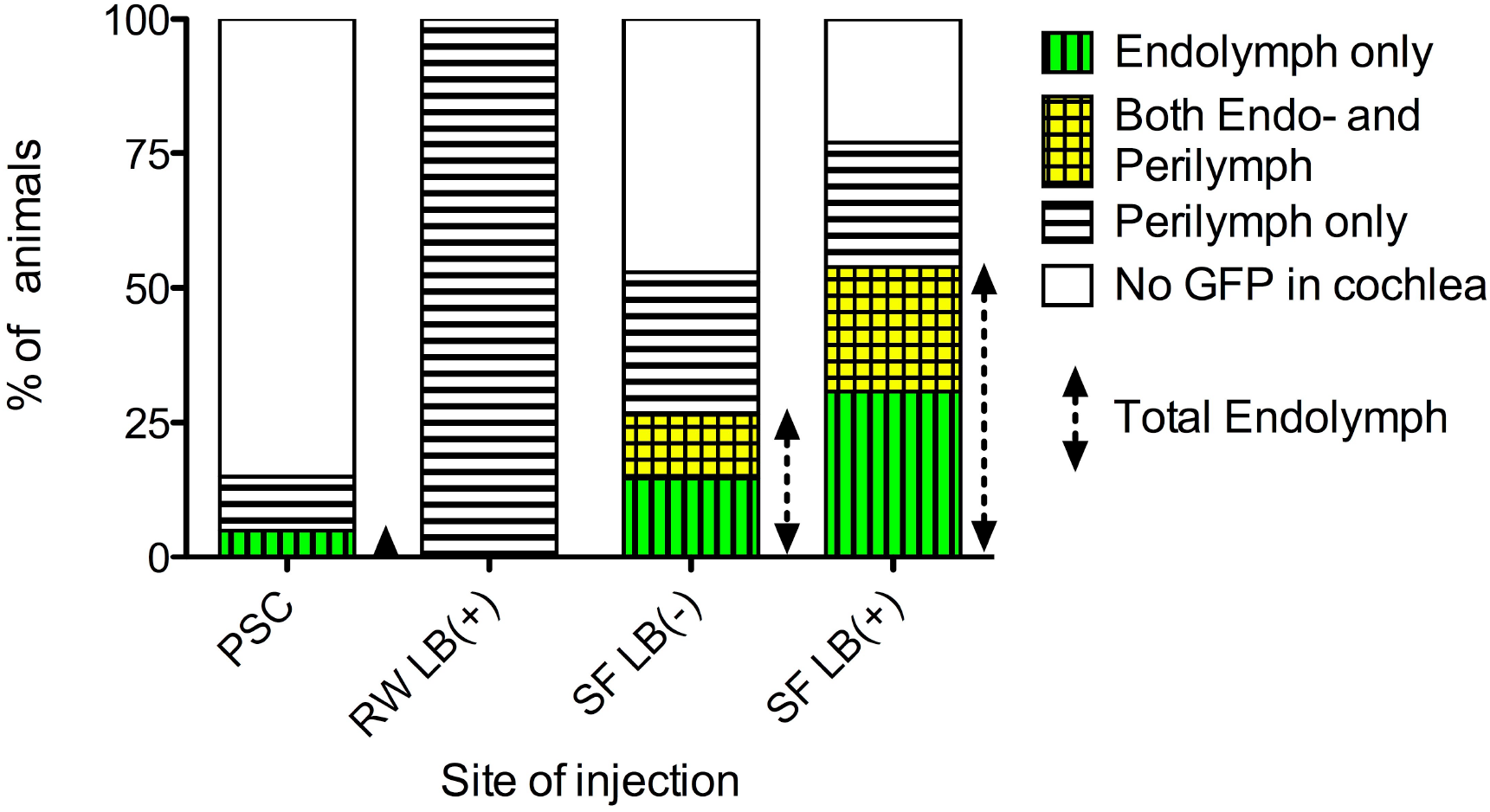
Localization of GFP-positive cells in the cochlea. Serial sections of injected animals were reviewed and tallied for the presence of GFP positive cells. Vertical bars indicate the percentage of injected animals with GFP-expressing cells adjacent to the endolymphatic fluid compartment alone, both the endolymphatic and perilymphatic fluid compartments, and the perilymphatic fluid compartment alone. In some animals, no GFP-expressing cells were identified in the cochlea suggesting that the virus had either leaked out from the injection site (all groups), or had not reached the cochlea (PSC group, see text). PSC= Posterior semicircular canal (n=20), RW= Round window (n=8), LB (+/-) indicates whether or not liquid bandage used to seal injection site. SF= Stylomastoid foramen (n= 34 LB-, n= 13 LB+)

### Posterior Semicircular Canal Injections (PSC)

In the virus-injected PSC group (n=14), 5.0% of mice had GFP-positive transgene expressing cells adjacent to the endolymphatic fluid compartment in the cochlea and 10% had GFP expression consistent with perilymphatic delivery. There were no positive cells in the cochlea in 85.0% of this group (Figure 2). In PSC animals without GFP expression in the cochlea, most GFP positive cells were localized in the vestibular labyrinth around the duct and ampulla of the posterior semicircular canal and the utricle (Figure 3). The mean threshold shift was 35 dB at 8 kHz when AdE-CMV-GFP was injected compared to 32 dB at 8 kHz when virus storage buffer was injected (n=8) (Figure 4). While both virus and storage buffer injections via the PSC approach caused a significant loss of hearing (p<0.001), they did not differ from each other (p>0.05). The complete set of statistical analyses is shown in the supplementary data table (Supplement 1).

**Figure 3.**
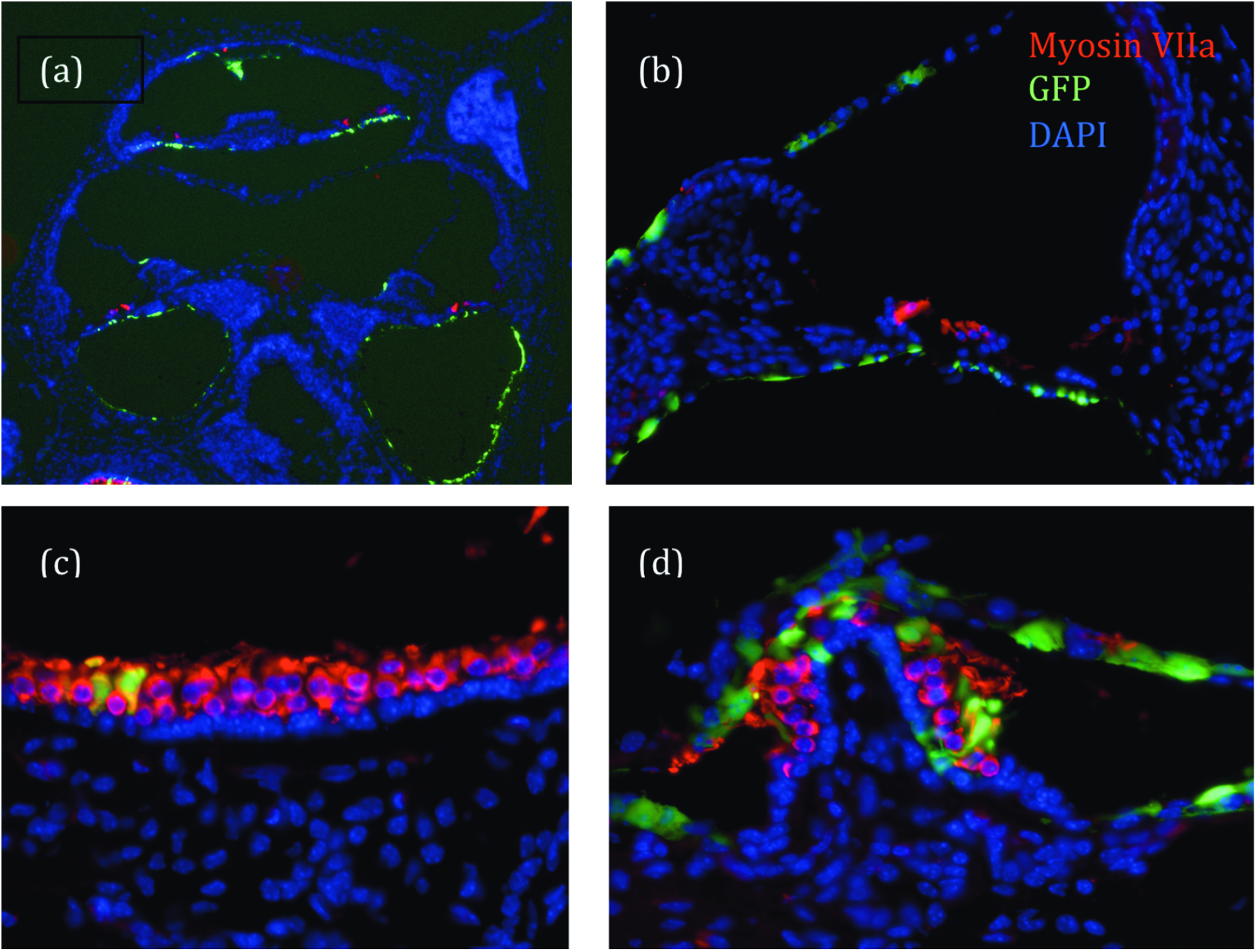
Posterior semicircular canal (PSC) injections. In most animals, there were no GFP-positive cells in the organ of Corti, but there was widespread expression in the vestibular labyrinth. **(a)** Overview of the cochlea. GFP-positive cells (green) adjacent to the perilymphatic space extending from the base to apex. **(b)** Higher magnification view showing Reissner’s membrane and the basilar membrane with GFP-expressing cells. Hair cells were stained with myosin VIIa (red). **(c)** There were GFP-positive cells in the endolymphatic and perilymphatic spaces of the utricle. In the vestibular sensory epithelia, the vast majority of GFP-expressing cells were supporting cells, not hair cells, based on nuclear position and absence of myosin VIIa labeling. **(d)** The ampulla showed a similar expression pattern as the utricle. GFP=green, myosin VIIa=red, DAPI=blue

**Figure 4.**
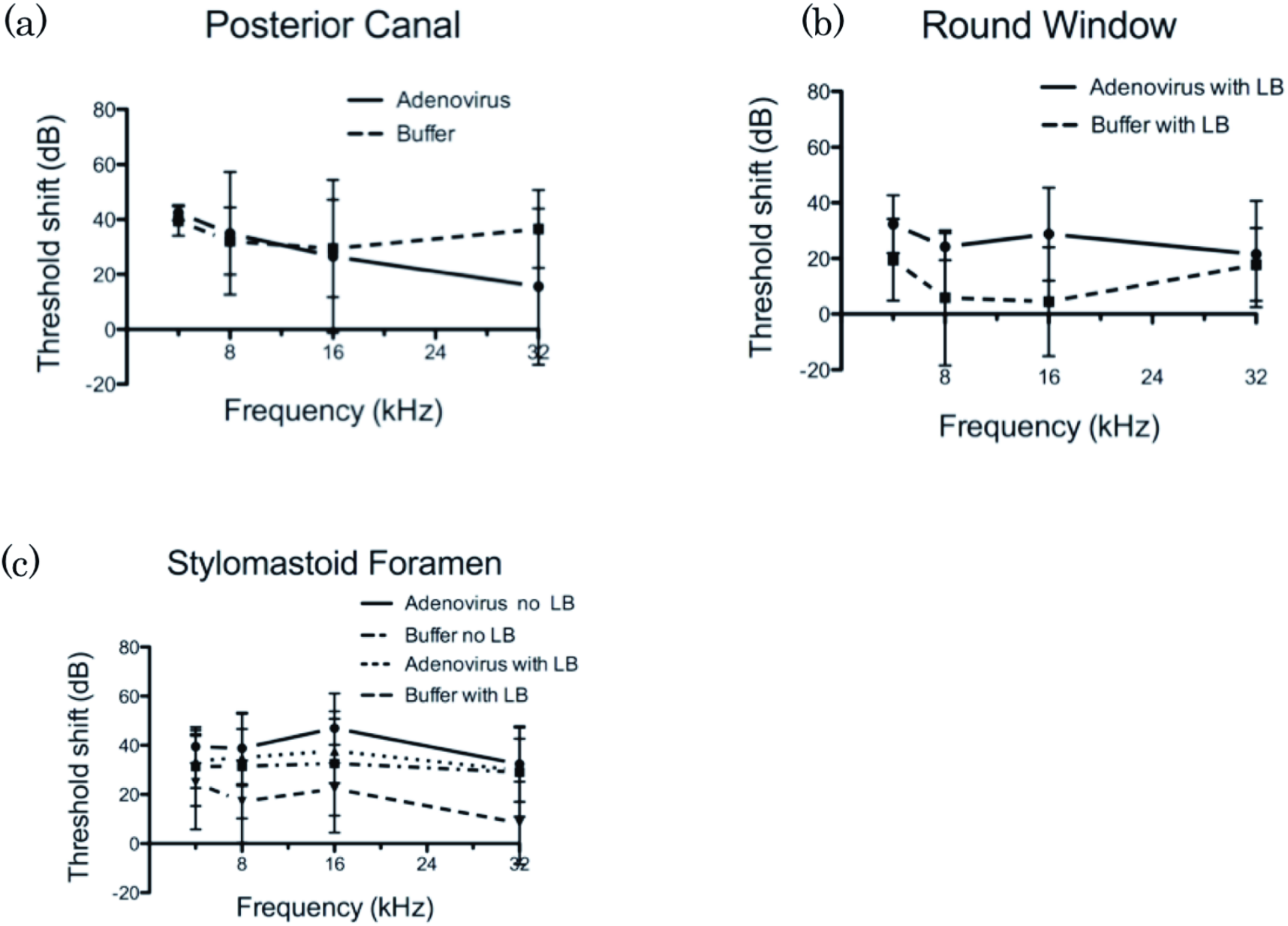
Impact of labyrinthine injections on hearing levels. ABRs are represented as shifts from averaged hearing thresholds in the same animals measured preoperatively. **(a)** <u>Posterior semicircular canal injections</u>. The hearing level of buffer and virus injected groups were almost the same. **(b)** <u>Round window injections</u>. When buffer alone was injected, there was minimal loss of hearing especially in the middle frequency range, however, when adenovirus was injected, mice showed a 20-35 dB threshold shift at each frequency. **(c)** <u>Stylomastoid Foramen injections</u>. Although the use of liquid bandage (LB) appeared to decrease the degree of hearing loss in both buffer and virus-injected animals with SF injections, this difference was not significant in our studies. For buffer injections without LB, threshold shifts were about 30-40 dB, but when the injection site was sealed with LB, the threshold shift was to 10-20 dB. For adenovirus injections using SF injection technique, with or without LB, the threshold shift was about 40dB.

### Round Window Injections (RW)

In the virus-injected RW group (n=7), all mice had GFP positive cells in the scala tympani from the basal turn through the middle turn. There were no GFP positive cells in the scala tympani of the apical turn, or in the scala vestibuli and the scala media of all turns (Figure 3). GFP positive cells were found in the epithelium lining the modiolus and the basilar membrane (Figure 5). The threshold shift was 24 dB at 8 kHz in the virus injected RW group and 6 dB at 8 kHz in the control loading buffer injected group (n=4) (Figure 4). The shift from preoperative hearing following a RW injection was significant for virus injected animals (p<0.01), but not for storage buffer (p>0.05).

**Figure 5.**
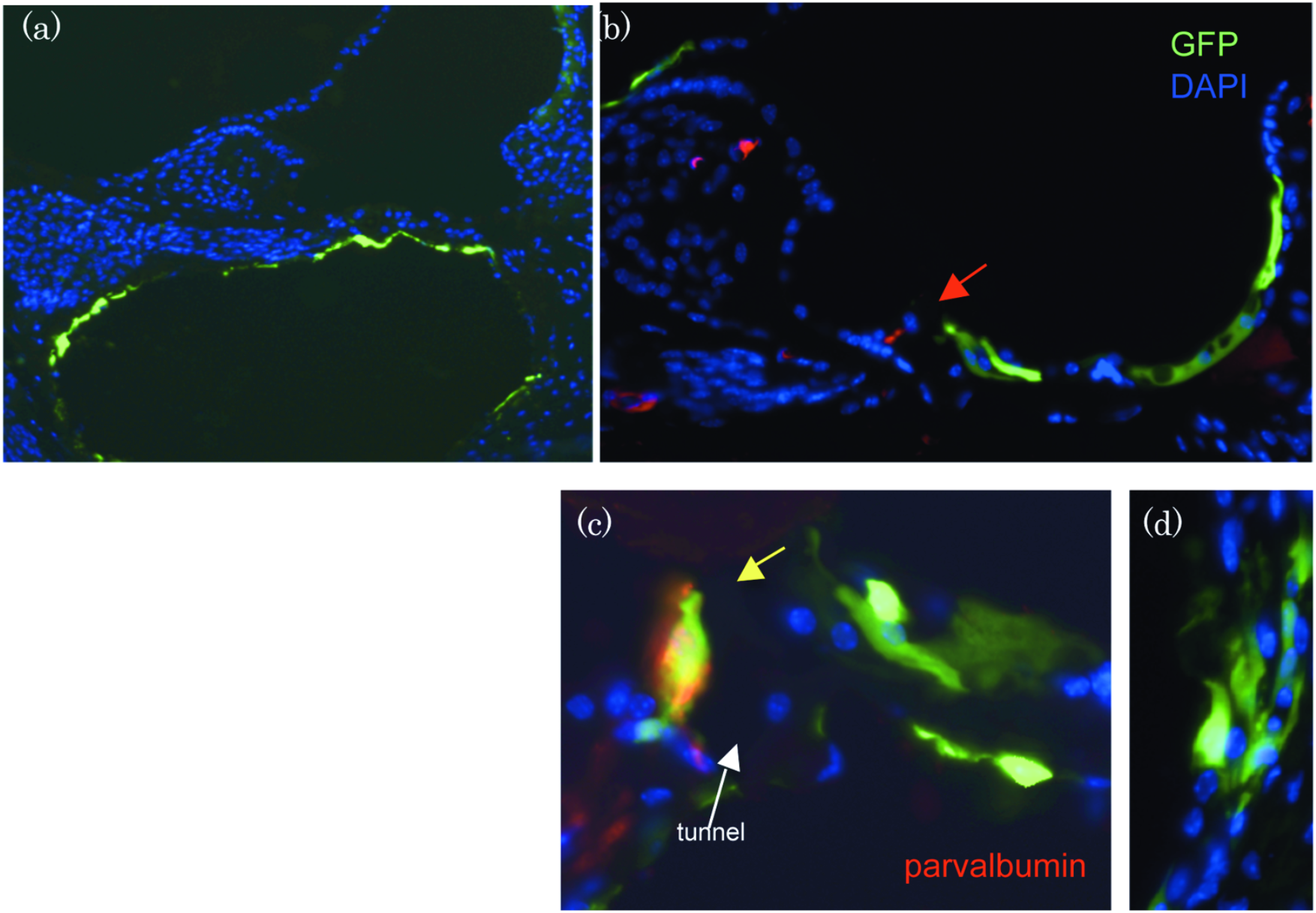
**(a)** Round window (RW) injections. In the RW group, all mice had GFP positive cells in the scala tympani from the basal turn through the second turn. **(b-d)** <u>Stylomastoid foramen (SF) injections.</u> Many GFP positive cells were found in the organ of Corti and Reissner’s membrane. GFP-expressing Deiters cells were identified in most animals. GFP positive Hensen cells or Claudius cells were often identified. In most animals, hair cells were unlabeled (red arrow in **b**). Rare, parvalbumin-GFP-double positive hair cells were also identified in some animals (yellow arrowhead in **c**). GFP-expressing cells were commonly found in the stria vascularis (d). GFP=green, parvalbumin=red, DAPI=blue. ST=scala tympani, SM=scala media, IHC=inner hair cell.

### Stylomastoid Foramen Cochleostomy Injections (SF)

In the virus-injected SF group (n=18), 26% of mice had GFP positive cells in the scala media when liquid bandage was not used (Figure 2). Without liquid bandage, the threshold shift was 39 dB at 8kHz in the virus injection group and 31 dB at 8kHz in the control loading buffer injected group (n=8). In contrast, when liquid bandage was used to seal the injection site (n=17), 54% of mice had GFP positive cells in the scala media (Figure 2). Based on location and morphology, the GFP expressing cells appear to be Deiters cells in most cases, although some Hensen’s cells and Claudius cells were also identified (Figure 5). For this group, the threshold shift was 35 dB at 8 kHz in the virus injection group and 17 dB at 8kHz in the control group (n=14) (Figure 4). In contrast to the RW approach, the hearing of SF buffer injected animals also differed from preoperative animals whether or not liquid bandage was utilized (p<0.05). Virus injections via the SF approach also caused a significant loss of hearing (p<0.001). We found that when using liquid bandage with the SF approach fewer animals had noticeable leakage during, or immediately after the procedure.

## 4. Discussion

Gene transfer studies in the cochlea are not limited to designing therapeutic strategies to alleviate auditory dysfunction, but can also contribute towards molecular genetic analysis of hearing. Mice provide an excellent model to study the genetics of hearing impairment because of the widespread availability of genetic models and newer ototoxicity protocols. Surgical access to the specific fluid compartments in the cochlea will be necessary for targeted therapeutic interventions. We analyzed the efficiency and hearing loss associated with several different surgical delivery routes to the mouse cochlea and found that injection via the stylomastoid foramen (SF) offers several advantages over previously described procedures to target the scala media.

An ideal route of virus delivery to the adult inner ear would be minimally invasive and non-destructive to hearing. The minute size of the inner ear, its encapsulation in dense bone and highly organized and differentiated structure place a number of restrictions on vector design and delivery. The mammalian inner ear comprises two distinct fluid spaces (endolymph, perilymph) that are essential for maintaining the electrochemical potential responsible for hearing sensitivity. The total volume of the inner ear fluid spaces range from less than two microliters in mouse to approximately two hundred microliters in the human (http://oto.wustl.edu/cochlea/). The endolymph and perilymph compartments are separated by thin membranes and surrounded by the temporal bone. Because of its delicate structure, surgical trauma to access the fluid spaces of the inner ear has a high risk of causing damage and hearing loss.

PSC injections were used to inject stem cells into the cochlea in a previous study and over 90% of transplanted cells were located in the perilymphatic space (Iguchi et al., 2004). The injected cells were assumed to go through the perilymphatic space of the vestibule. In our present study, viral vectors seemed to follow the same route. In most cases, GFP positive cells were located around the ampulla of posterior semicircular canal and the utricle. Some mice had GFP positive cells inside the utricle, but only a small number of mice showed GFP positive cells in the cochlea. Although the ductus reuniens interconnects the endolymphatic fluid spaces between the vestibular system and the cochlea, it appears to function as a check valve. For these reasons, we were not able to consistently and selectively access the endolymphatic fluid compartment in the cochlea via the semicircular canal approach. We hypothesize that when elevated injection pressures breach the endolymphatic compartment, virus is able to penetrate through holes in the membranous labyrinth into the endolymph. If the injection speed or volume is increased, the number of mice with GFP positive cells in the scala media would likely increase, but at the expense of postoperative hearing level. Even under our relatively mild conditions, hearing preservation was already compromised by the procedure.

Several groups have used microinjection via round window into the scala tympani to deliver viral vectors to the inner ear (Raphael et al., 1996). This method directly accesses the scala tympani and can consequently cause some hearing loss and imbalance from traumatic disruption of the inner ear. In the current study, the hearing threshold shift was only 6 dB at 8kHz in the control group. The use of liquid bandage sealed the injection site and prevented significant perilymph leakage from the round window. Injection of adenovirus did cause a significant change in hearing compared to control injections. In our studies using this approach, all GFP positive cells were located in the scala tympani. Even though there were robust GFP positive cells in the epithelia bordering the modiolus and the basilar membrane, there were no GFP positive cells in the organ of Corti, the scala media or within the modiolus itself. Therefore, it appears that adenovirus is unable to penetrate the basilar membrane or a bony wall of the modiolus. This is consistent with the result in previous report (Venail et al., 2007). In contrast, when adeno-associated virus (AAV) is injected via the RW, transduced cells are found within all fluid compartments (Liu et al., 2007). We have also found that AAV-CMV-GFP injected via the PSC can target cells in both endolymphatic and perilymphatic fluid compartments (unpublished). The differences between Adenovirus and AAV may be due to the size of the capsid (∼25 nm in diameter) or effects of virus binding to competing receptors (Excoffon et al., 2006; Venail et al., 2007)

In addition to these two methods, we have developed a new approach, via the stylomastoid foramen, for gene delivery into the mouse scala media. This approach has the advantage of increasing targeting to the organ of Corti, while preserving some hearing. The stylomastoid foramen is located between the round window and the oval window where only a thin layer of bone overlies the scala media. Therefore, when the cochlear duct is punctured, the endolymphatic and perilymphatic spaces do not become connected and can maintain their potential difference. Because the buffers in our study contained a visible blue dye, we were able to directly monitor leakage and delivery to the cochlea during each injection. To minimize endolymph leakage and loss of virus following injection, we found that liquid bandage was critical. Although the hearing levels were not significantly better using liquid bandage, the percentage of mice that had GFP positive cells in the scala media was 54% in the SF group with liquid bandage compared to 26% in the SF group without liquid bandage. This corresponded to our observation that liquid bandage decreased the amount of leakage from the injection site, during and after the procedure.

Early studies with first and second-generation adenoviral vectors showed some toxicity related to expression of viral proteins and a host inflammatory response. Subsequent studies using advanced generation adenoviral vectors showed less toxicity and supported the potential use of adenovirus in inner ear gene therapy model systems (Holt et al., 1999; Luebke et al., 2001; Luebke et al., 2001; Holt, 2002; Wenzel et al., 2007). Even with advanced generation adenovirus we found that virus injections compared to buffer injections had adverse affects on hearing regardless of the route of delivery. This effect was less dramatic, but still significant for injections via the RW and PSC, compared to the SF. As described by others previously, this suggests that adenoviral transduction, either directly, or indirectly through an immune response may have the undesirable effect of causing tissue damage resulting in hearing loss (Lalwani and Mhatre, 2003; Nemerow, 2009). This toxicity could potentially be decreased through use of helper dependent adenovirus (Wenzel et al., 2007), or pretreatment with anti-inflammatory agents (Seregin et al., 2009). Although there may be direct toxicity of adenovirus-receptor interactions in the absence of viral gene expression, some experiments in guinea pig using the same adenovirus backbone suggest that this effect is minimal (Luebke et al., 2001; Luebke et al., 2001; Nemerow, 2009). Compared to guinea pig, the smaller size of the mouse cochlea or its immune response may make it more sensitive to these effects.

## 5. Conclusion

We have developed a new method for targeted delivery to the scala media of the mouse inner ear that preserves some hearing. When adenovirus is injected via the round window, transduced cells are found bordering only the scala tympani. Adenovirus vector seems unable to penetrate the basilar membrane or the bony wall of the modiolus. Delivery to the cochlea via the semicircular canals is also not very efficient. In contrast, our new method, via a stylomastoid foramen cochleostomy, increases the likelihood of adenovirus gene transfer to the scala media and cells in the organ of Corti and stria vascularis. We also describe the use of liquid bandage to minimize of endolymphatic fluid during this type of delivery and increase the success of targeting the organ of Corti. Our technique is a useful addition to existing methods when targets of viral or small molecule delivery are in the scala media. In contrast, adenoviral delivery via the semicircular canals is a more efficient route to target the vestibular labyrinth while round window injections more efficiently target the cochlear perilymphatic compartment. The advanced generation adenovirus we have used in our studies cause some hearing loss in contrast to gutted adenovirus and AAV, however, these vectors have the advantage of more rapid construction of high titer molecular variants to allow for in vivo screening.

The ability to target delivery of adenovirus using several routes of injection will facilitate in vivo biological testing of candidate molecules implicated in multiple aspects of inner ear physiology and regeneration.

## Acknowledgements

Thanks to Jeffrey Chamberlain (University of Washington) and Andrea Amalfitano (Michigan State University) for C7-HEK 293 cells, backbone vectors and advice on preparation of replication defective E1-/E3-/preterminal protein-/polymerase-adenovirus.

